# Validity of Two-Dimensional Static Footprint in Medial Longitudinal Arch Evaluation and the Characteristics of Athletes’ Footprints

**DOI:** 10.1101/2020.02.10.941633

**Authors:** Lingli Zhang, Dali Yu, Le Lei, Yuanwu Gao, Junjie Dong, Zhusheng Yu, Yu Yuan

**Author notes:** Correspondence and requests for materials should be addressed to: Dr. Yu Yuan, Address: School of Physical Education & Sports Science, South China Normal University, 55 Zhongshan Road West, Tianhe District, Guangzhou, 510631, China, Dr. Zhusheng Yu, Address: Department of Kinesiology, Shanghai University of Sport, 188 Hengren Road, Yangpu District, Shanghai, 200438, China.

## Abstract

**Background:** We aimed to explore the validity of two-dimensional static footprint analysis in medial longitudinal arch evaluation as well as the characteristics of athletes’ footprints to provide a basis for the evaluation and selection of athletes.

**Methods:** Experiment One: Twenty-nine high level athletes (runners and jumpers) and forty normal college students were selected. Based on the X-ray photos taken of the medial foot, we measured the calcaneal inclination angle, the calcaneal–first metatarsal angle and the ratio of height to length of the medial longitudinal arch. We collected indicators of two-dimensional static footprints. Experiment Two: 106 high level athletes (runners and jumpers) and 104 normal college students were selected. We also collected indicators of two-dimensional static footprints.

**Results:** The average measuring the Interclass Correlation Efficient (ICC) of calcaneal inclination angle, calcaneal–first metatarsal angle, the ratio of height to length of the medial longitudinal arch, the width of ball, arch and heel, the length of footprint and each toe, Chippaux-Smirak Index (CSI) and Staheli Index (SAI) were higher than 0.800. Regardless of athletes or college students, male or female, the correlation between CSI, SAI and calcaneal inclination angle, calcaneal–first metatarsal angle, the ratio of height to length of the medial longitudinal arch was statistically significant *(P<0.05)*. College students’ CSI of the right foot is significantly higher than that of the left foot regardless of gender *(P<0.05)*.

**Conclusions:** We prove the qualification of CSI and SAI in medial longitudinal arch evaluation and explain that the relative height of medial longitudinal arch is an important indicator in track and field.

## Introduction

The analysis of two-dimensional footprint is important in many fields such as biomechanics, ergonomics, forensic medicine and anatomy. The research results in this field have been widely used in crime investigation, clinical diagnosis, footwear design and the selection of athletes^1^. However, there is no uniform standard for footprint measurement and evaluation.

Several methods have been adopted in traditional arch measurement including ink^2^, footprint mat^3^, inkless paper system^1,4^, etc. However, footprints collected in the traditional ways lack a distinct outline as well as durability so that they are difficult to be verified after long-term storage. The Yushi Footprint Measurer obtained a China national invention patent of independent intellectual property right for an arch measuring method and device in 2009. This study applied the Yushi Footprint Measurer to measure the footprint (Invention patent number: ZL200910046464.5). The footprints collected in this way are clearly visible, and they can be permanently preserved and repeatedly verified.

The medial longitudinal arch is one of the most important variables in studying footprint, and the collapse of the medial longitudinal arch is a sign of flatfoot^5,6^. Flatfoot is a common morphological variation of the foot affected by genetic inheritance, gender, obesity, footwear and other factors^6,7^. Flatfoot influences physical activities^8^ so it is important to diagnose and identify flatfoot scientifically. Compared with radiology and ultrasonography, footprint analysis is less expensive, faster, more effective and easier to perform^8–10^. Two-dimensional static footprint analysis is one of the commonly used methods in medial longitudinal arch evaluation. However, there is no unified standard for the acquisition and evaluation of footprint. The purpose of the first part in this study was to verify the accuracy of “Two-Dimensional Static Footprint in Medial Longitudinal Arch Evaluation” as a methodology, and the second part was to compare the differences between athletes of high level and normal university students with Two-Dimensional Static Footprint. The purpose of this study was to explore the validity of two-dimensional static footprint analysis in medial longitudinal arch evaluation as well as the characteristics of athletes’ footprints to provide a basis for the evaluation and selection of athletes.

## Materials and Methods

### Ethics statement

Ethical approval was obtained from the Ethics Committee of the Shanghai University of Sport Human Subjects Research Review Committee (Shanghai, China, Approval Number: 2014029). The study followed the protocols approved by the committee, and the subjects’ confidentiality was strictly maintained throughout the study.

### Experiment One

#### Participants

Twenty-nine students (12 males and 17 females) from Xinzhuang training base of Shanghai Physical Education Technology Institute and School of Competitive Sports affiliated to Shanghai University of Sport were selected as subjects of sport group from November 3 to December 24, 2014. They were high level runners and jumpers. We chose 40 college students (16 males and 24 females) who were healthy without apparent malformation of foot and majored in academic science.

Firstly, we contacted their coaches or teachers, and explained the purpose, procedure and relative points for attention to the athletes or students to ensure that the subjects could cooperate to complete the test actively and voluntarily. Informed consent was read and signed by each subject before the test.

#### Methods of Measurement

We have done the following to minimize the effect of the testers as well as measuring apparatuses on testing results. Each of the measuring apparatus had been calibrated before the test. During the test, the measuring of height and weight was done by the same tester, the collection of the footprint of the left foot of all subjects was done by the same tester, the collection of the footprint of the right foot of all subjects was done by the same tester, and the analysis of the indicators of the footprints was done by two testers. All the testers accepted professional training before the test. The X-ray photographer is professional with certificate.

#### Information of the subjects

We used a questionnaire to ask for the subjects’ age, sport-specialization and level. A variety of anthropometric parameters, including weight and height, were measured while the subjects were in light clothing without shoes. Weight was measured with the weighing scale to the nearest 0.01kg and height was measured with a hypsometer to the nearest 0.01cm. The data of the subjects including age, height and weight are in Table 1. Height and weight were tested by the trained tester.

**Table 1.** Baseline information of the subjects

| Group | Sex | Experiment One |  |  |  | Experiment Two |  |  |  |
| --- | --- | --- | --- | --- | --- | --- | --- | --- | --- |
|  |  | N | Age<br>(year) | Height<br>(cm) | Weight<br>(kg) | N | Age<br>(year) | Height<br>(cm) | Weight<br>(kg) |
| Control | Male | 16 | 24.19±2.11 | 173.47±5.28 | 72.70±9.15 | 54 | 21.17±2.16 | 175.34±5.25 | 65.51±7.04 |
|  | Female | 24 | 23.17±1.44 | 165.59±4.79 | 56.23±6.48 | 50 | 20.84±1.92 | 164.45±4.28 | 53.85±9.69 |
| Sport | Male | 12 | 20.25±2.90 | 180.89±3.71 | 71.61±6.77 | 56 | 21.82±3.58 | 177.91±6.22 | 67.46±9.64 |
|  | Female | 17 | 20.24±2.49 | 167.88±6.22 | 56.31±5.43 | 50 | 21.40±2.60 | 166.25±5.37 | 54.92±6.79 |

#### Taking X-ray photos of medial foot

We used N400 digital medical X-ray photography system (Neusoft Co., China) to take X-ray photos of medial foot. N400 applies portable flat panel detector and its digital system takes X-ray photos and saves them in digital format automatically. It can be applied to any part of the body. The X-ray output is of high accuracy with the minimum dose. The X-ray photographer is professional with certificate. We required that during the test of the X-ray, the subjects should stand naturally with feet shoulder-width apart, middle toes of both feet pointing to the front. The subjects were asked to stand on a horizontal stool during the test of the X-ray.

When the subjects stood firm, the photographer focused the X-ray on the medial foot of the subjects with the tube current of 100 milliampere (mA), the tube voltage of 60 kilovolt (KV), the detector distance of 1800 millimeter (mm) and the time of exposure of 0.16 second. After the X-ray photos of bilateral medial feet were taken, we exported the photos and measured them with Dicom Viewer Medical Image Tool 1.0 in computer. In order to test the validity of the testers, two testers measured the following indicators:

1. Calcaneal inclination angle (Figure 1) Calcaneal inclination angle is the inclination of the tangent of the lower edge of calcaneal to the plane on which the foot stands^11^.
2. Calcaneal-first metatarsal angle Calcaneal-first metatarsal angle is also called medial longitudinal arch angle, that is the inclination of the line joining the bottom of the first metatarsal and the bottom of talonavicular joint to the line joining the bottom of calcaneal and the bottom of talonavicular joint^12^.
3. Ratio of height to length of the medial longitudinal arch (Figure 1) The length of the medial longitudinal arch is the length of the line joining the bottom of calcaneal and the bottom of the first metatarsal. The height of the medial longitudinal arch is the vertical distance between the bottom of talonavicular joint and the line joining the bottom of calcaneal and the bottom of the first metatarsal. As one of the commonly used methods in arch evaluation, the ratio of height to length of the medial longitudinal arch is more accurate and reliable than the comparison of the height of the medial longitudinal arch alone.

**Figure 1.**
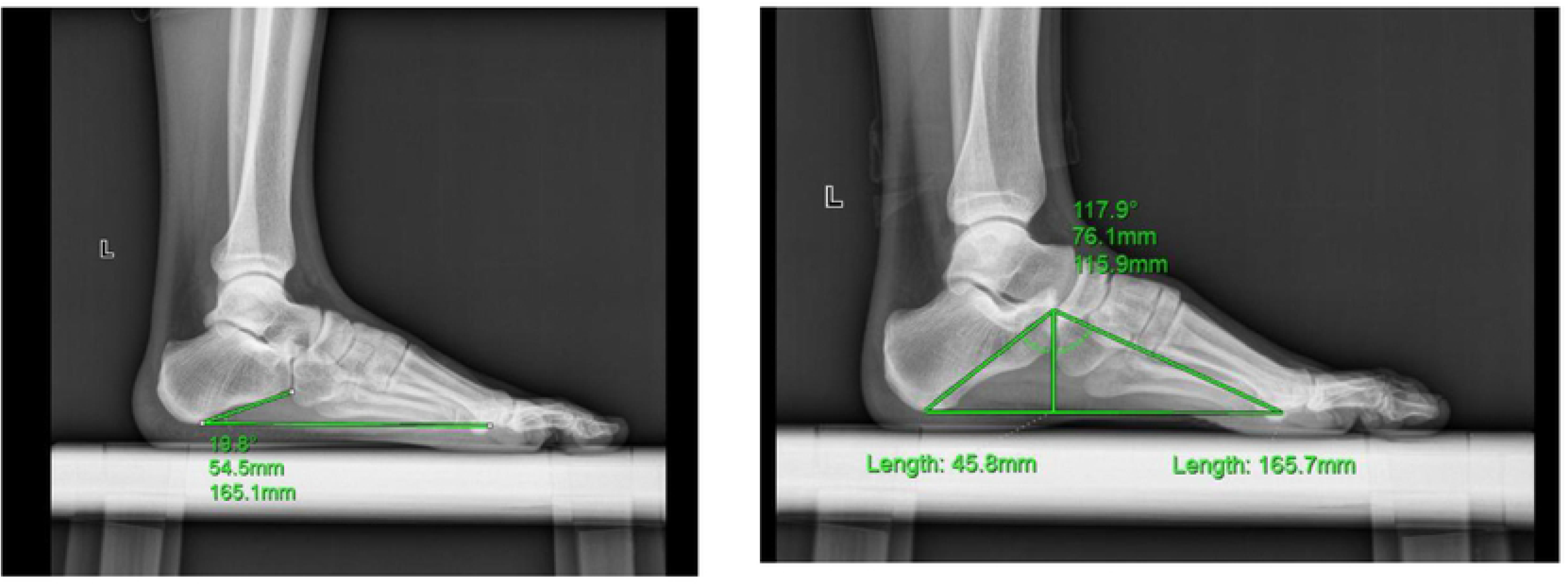
The indicators of X-ray photos of calcaneal inclination angle and medial foot evaluating medial longitudinal arch.

#### Collecting two-dimensional static footprints

The dipping appliance for footprint (L35cm×W25cm), an important part of the Yushi Footprint Measurer, obtained a patent for the invention, and is an independent intellectual property right of the People’s Republic of China (Invention patent number: ZL200910046464.5), was used to collect the footprints. The following consumable materials were used in this experiment: a red or blue inkpad box, 50g of ink, A4 papers, cleaning agent and coarse paper.

Firstly, the footprint measurer was put on the floor horizontally. The subjects then dipped the inkpad with barefoot and stood on the A4 paper. We required that during the test of two-dimensional static footprints, the subjects should stand naturally with feet shoulder-width apart, middle toes of both feet pointing to the front. The subjects were asked to stand steadily that no movement of the foot was permitted until the clear footprint was formed. The footprints of left and right foot were tested by two trained testers. The footprints collected in this way endure long-term preservation and can be repeatedly verified.

In order to test the validity of the testers, after the collection of bilateral static footprints of all the subjects, we copied the footprint with the copying machine and each of the two testers measured one copy separately. Two testers divided footprint into the toe zone, the sole zone, the arch zone and the heel zone, drew the inner tangent and outer tangent of the footprints according to Reel^4^ and measured the following indicators (Figure 2):

1. Ball Width: distance between the outermost point of the fifth toe joint and the innermost point of the metatarsal bone
2. Arch Width: width of the narrowest area of the arch zone
3. Heel Width: distance between the innermost concave of the calcaneal and the outermost point of the calcaneal
4. Foot Length: distance between the rear edge of the heel and the front edge of the toes
5. Toe Length: the maximum length of the toes
6. Chippaux-Smirak Index (CSI) = (Arch Width/ Ball Width) × 100%
7. Staheli Arch Index (SAI) = (Arch Width/ Heel Width) × 100%

**Figure 2.**
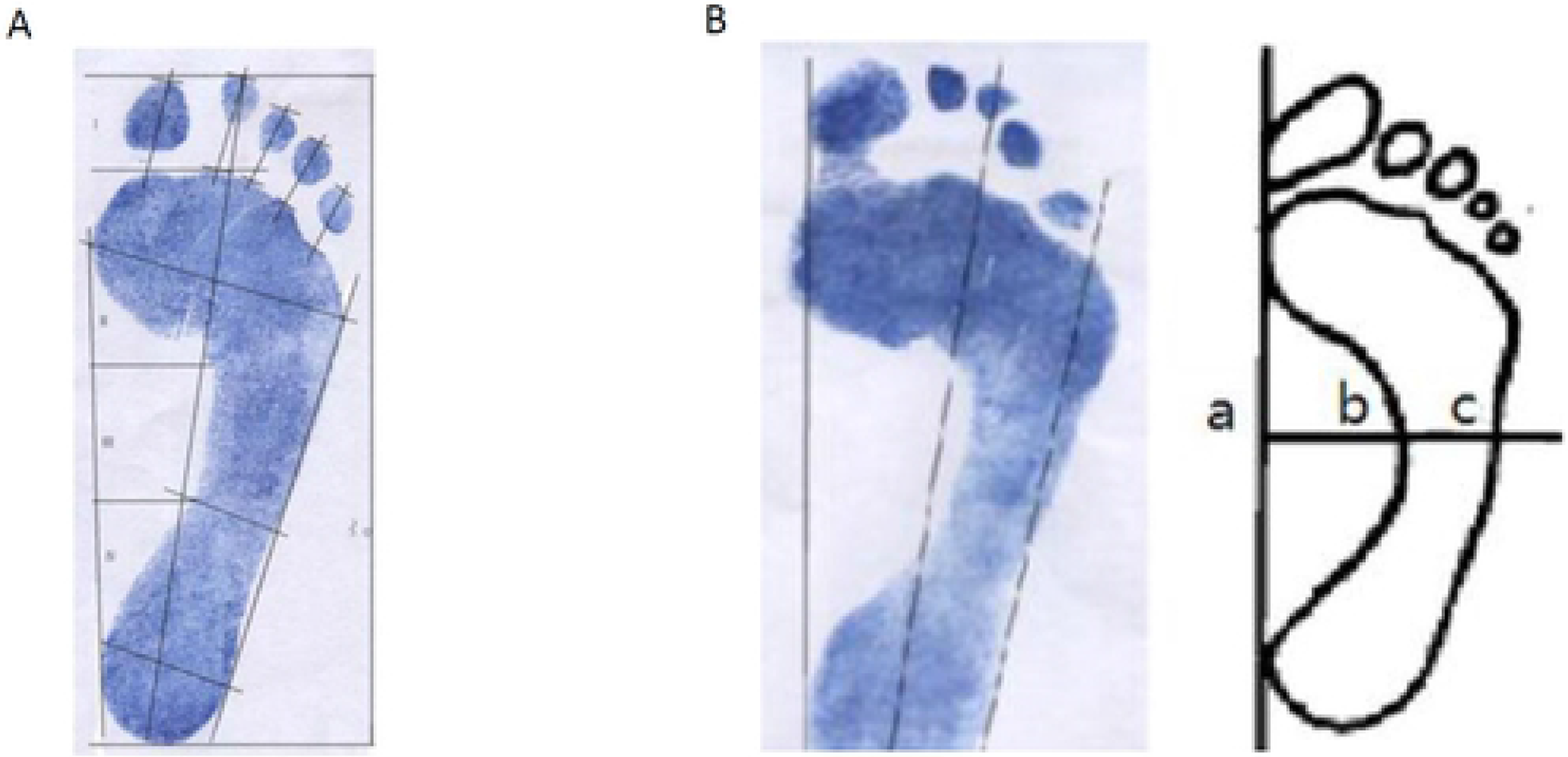
A The indicators of two-dimensional static footprint; B “Three Lines Method” (left) and the “Ratio Method” (right).

### Experiment Two

#### Participants

We selected 106 athletes of high level, who are specialized in running and jumping (56 males and 50 females), including 85 national level-1 athletes and 18 master sportsmen, from Xinzhuang training base of Shanghai Physical Education Technology Institute, School of Competitive Sports affiliated to Shanghai University of Sport, East China Normal University, and Shanghai professional sports teams from April 14, 2015 to March 29, 2016. We also selected 104 normal university students (54 males and 50 females) of similar age and height with the selected athletes from Department of Kinesiology from Shanghai University of Sport, who are majored in academic science with no specialization in any sport. All subjects were healthy without apparent malformation of foot. Informed consent was read and signed by each subject before the test to ensure that the subjects could complete the test actively and voluntarily.

### Methods of Measurement

#### Information of the subjects

The data of the subjects’ information including age, height and weight are recorded in Table 1. For athletes, we also record the sports in which they are specialized and their level.

#### Collecting two-dimensional static footprints

The method is the same as that in Experiment One.

### Statistical analysis

SPSS 22.0 software (SPSS, Chicago, IL, USA) was used to analyze the data, which were expressed as mean ± standard deviation (SD). The Interclass Correlation Efficient (ICC) was applied to analyze the reliability of two testers who measured the indicators of two-dimensional static footprints as well as X-ray photos by data evaluation through average measuring ICC^13^. The Pearson correlation was adopted to analyze the association between indicators of two-dimensional static footprints and indicators of X-ray photos of medial foot. Mean values of two testers’ measuring data are accepted as the final data. Statistical differences were calculated with independent T-test and paired samples T-test. P values < 0.05 were considered statistically significant.

## Results

### Reliability of the testers

In order to ensure the reliability of the two testers who measured the indicators of two-dimensional static footprints as well as X-ray photos, they did not take part in previous steps of the experiment. They read the evaluating method of two-dimensional static footprint carefully and measured the indicators of the footprints in Experiment One. Mean values of two testers’ measuring data are accepted as the final data. The Interclass Correlation Efficient (ICC) was used to analyze the data through average measuring ICC. In Table 2, average measuring ICC of calcaneal inclination angle, calcaneal–first metatarsal angle, the ratio of height to length of arch, the width of ball, arch and heel, the length of footprint and each toe, CSI and SAI were higher than 0.900 (*ICC>0.800*), indicating that all the indicators are of high reliability.

**Table 2.** The reliability of the testers measuring indicators of footprints and X-ray photos of medial foot

| Indicators | Average measuring ICC |
| --- | --- |
| Calcaneal Inclination Angle | 0.943* |
| Calcaneal–First Metatarsal Angle | 0.966* |
| Ratio of Height to Length of Arch | 0.957* |
| Ball Width | 0.960* |
| Arch Width | 0.981* |
| Heel Width | 0.951* |
| Foot Length | 0.917* |
| CSI | 0.976* |
| SAI | 0.975* |
| Length of Great Toe | 0.933* |
| Length of the Second Toe | 0.960* |
| Length of the Third Toe | 0.959* |
| Length of the Fourth Toe | 0.962* |
| Length of the Fifth Toe | 0.954* |

\* $ICC > 0.800$ : The reliability of the testers is good.

### The correlation between CSI, SAI and calcaneal inclination angle, calcaneal–first metatarsal angle, the ratio of arch height to arch length

Calcaneal inclination angle, calcaneal-first metatarsal angle and the ratio of arch height to arch length are direct indicators in medial longitudinal arch evaluation. The Pearson correlation was used to analyze the association between CSI and SAI in data analysis of two-dimensional static footprints and X-ray photos of medial foot. Table 3 shows that the correlation between CSI, SAI and calcaneal inclination angle, calcaneal-first metatarsal angle and the ratio of arch height to arch length is statistically significant regardless of gender or whether they are athletes or not *(P<0.05)*.

**Table 3.** The correlation between CSI, SAI and the indicators of X-ray photos

| Group | Sex | Indicators | CSI |  | SAI |  |
| --- | --- | --- | --- | --- | --- | --- |
|  |  |  | Left foot | Right foot | Left foot | Right foot |
|  |  |  | r | r | r | r |
| Control | Male<br>(n=16) | Calcaneal Inclination Angle (°) | -0.572* | -0.705** | -0.562* | -0.726** |
|  |  | Calcaneal–First Metatarsal Angle (°) | 0.783** | 0.864** | 0.742** | 0.853** |
|  |  | Ratio of Height to Length of Arch | -0.752** | -0.874** | -0.708** | -0.857** |
|  | Female<br>(n=24) | Calcaneal Inclination Angle (°) | -0.652** | -0.695** | -0.583** | -0.661** |
|  |  | Calcaneal–First Metatarsal Angle (°) | 0.681** | 0.711** | 0.620** | 0.722** |
|  |  | Ratio of Height to Length of Arch | -0.628** | -0.689** | -0.560** | -0.673** |
| Sport | Male<br>(n=12) | Calcaneal Inclination Angle (°) | -0.713** | -0.741** | -0.745** | -0.719** |
|  |  | Calcaneal–First Metatarsal Angle (°) | 0.751** | 0.730** | 0.762** | 0.721** |
|  |  | Ratio of Height to Length of Arch | -0.707* | -0.741** | -0.713** | -0.728** |
|  | Female<br>(n=17) | Calcaneal Inclination Angle (°) | -0.593* | -0.562* | -0.575* | -0.546* |
|  |  | Calcaneal–First Metatarsal Angle (°) | 0.648** | 0.538* | 0.654** | 0.554* |
|  |  | Ratio of Height to Length of Arch | -0.658** | -0.576* | -0.675** | -0.546* |

\*. $P < 0.05$ , \*\*. $P < 0.01$ .

Table 3 presents CSI and SAI of male college students was moderately correlated to calcaneal inclination angle of left foot, other indicators had a high correlation with CSI and SAI of male college students and athletes of high level. Table 3 also shows that CSI and SAI of female college students were highly correlated to calcaneal–first metatarsal angle of right foot, other indicators had a moderate correlation with CSI and SAI of female college students and athletes of high level. The trend of CSI and SAI were the same.

### Comparison of the indicators of two-dimensional static footprints between athletes and college students

CSI differences between male athletes and male college students are not statistically significant. Female athletes’ mean CSI is significantly smaller than that of female college students in Table 4 *(P<0.05)*.

**Table 4.**
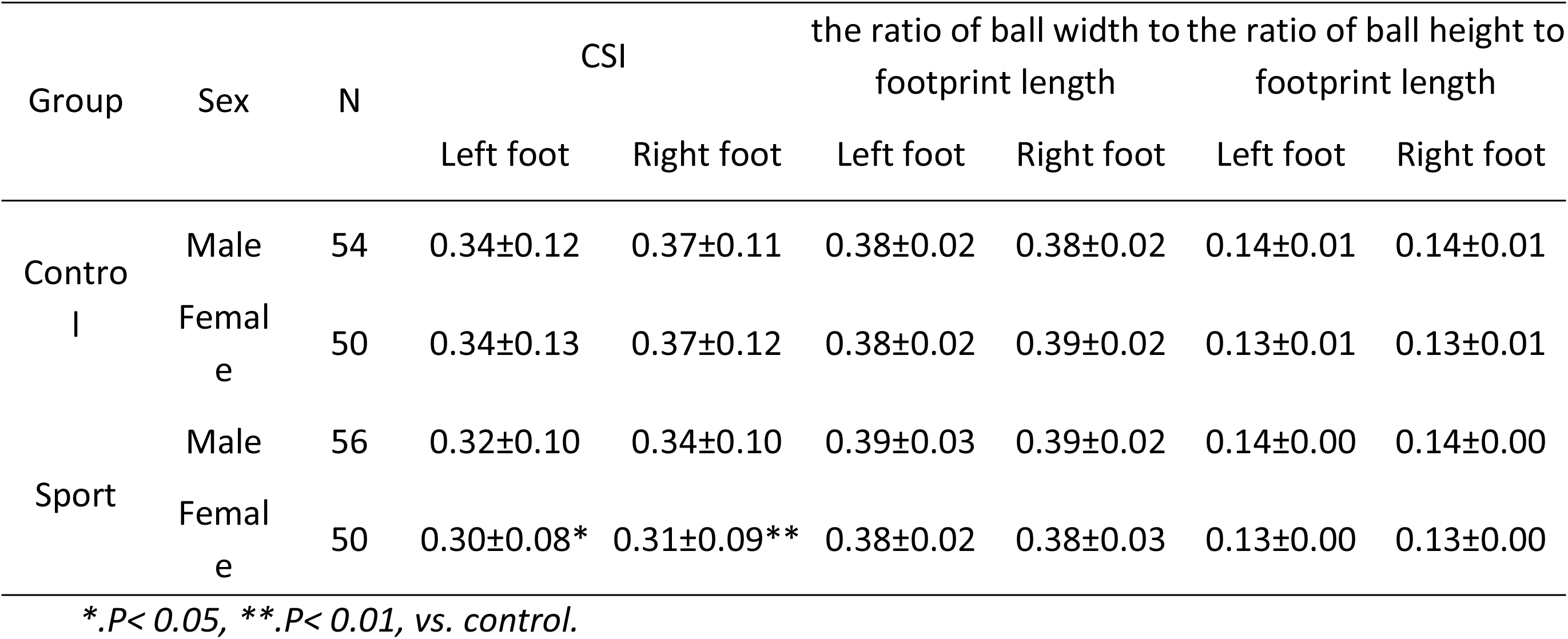
Comparison of CSI, the ratio of ball width and height to footprint length between athletes and college students

Table 4 and table 5 show that regardless of gender, there are no significant difference in the ratio of toe length to footprint length, the ratio of ball width to footprint length and the ratio of footprint length to body height between athletes and college students. Table 6 shows that college students’ CSI of the right foot is significantly higher than that of the left foot *(P<0.05)*. However, there is no similar bilateral CSI differences in athletes.

**Table 5.** Comparison of the ratio of toe length to footprint length between athletes and college students

| Group | Sex | N | Ratio | Left foot | Right foot |
| --- | --- | --- | --- | --- | --- |
| Control | Male | 54 | Length of Great Toe | 0.16±0.01 | 0.17±0.01 |
|  |  |  | Length of the Second Toe | 0.15±0.01 | 0.15±0.02 |
|  |  |  | Length of the Third Toe | 0.12±0.02 | 0.13±0.01 |
|  |  |  | Length of the Fourth Toe | 0.11±0.02 | 0.11±0.01 |
|  |  |  | Length of the Fifth Toe | 0.09±0.01 | 0.09±0.01 |
|  | Female | 50 | Length of Great Toe | 0.17±0.01 | 0.16±0.02 |
|  |  |  | Length of the Second Toe | 0.14±0.01 | 0.14±0.01 |
|  |  |  | Length of the Third Toe | 0.12±0.02 | 0.12±0.01 |
|  |  |  | Length of the Fourth Toe | 0.10±0.02 | 0.10±0.01 |
|  |  |  | Length of the Fifth Toe | 0.09±0.01 | 0.09±0.01 |
| Sport | Male | 56 | Length of Great Toe | 0.16±0.02 | 0.16±0.02 |
|  |  |  | Length of the Second Toe | 0.15±0.01 | 0.15±0.01 |
|  |  |  | Length of the Third Toe | 0.12±0.02 | 0.12±0.02 |
|  |  |  | Length of the Fourth Toe | 0.11±0.02 | 0.10±0.02 |
|  |  |  | Length of the Fifth Toe | 0.09±0.02 | 0.09±0.01 |
|  | Female | 50 | Length of Great Toe | 0.16±0.02 | 0.16±0.01 |
|  |  |  | Length of the Second Toe | 0.15±0.01 | 0.14±0.02 |
|  |  |  | Length of the Third Toe | 0.12±0.02 | 0.11±0.02 |
|  |  |  | Length of the Fourth Toe | 0.11±0.02 | 0.10±0.02 |
|  |  |  | Length of the Fifth Toe | 0.09±0.01 | 0.09±0.01 |

**Table 6.**
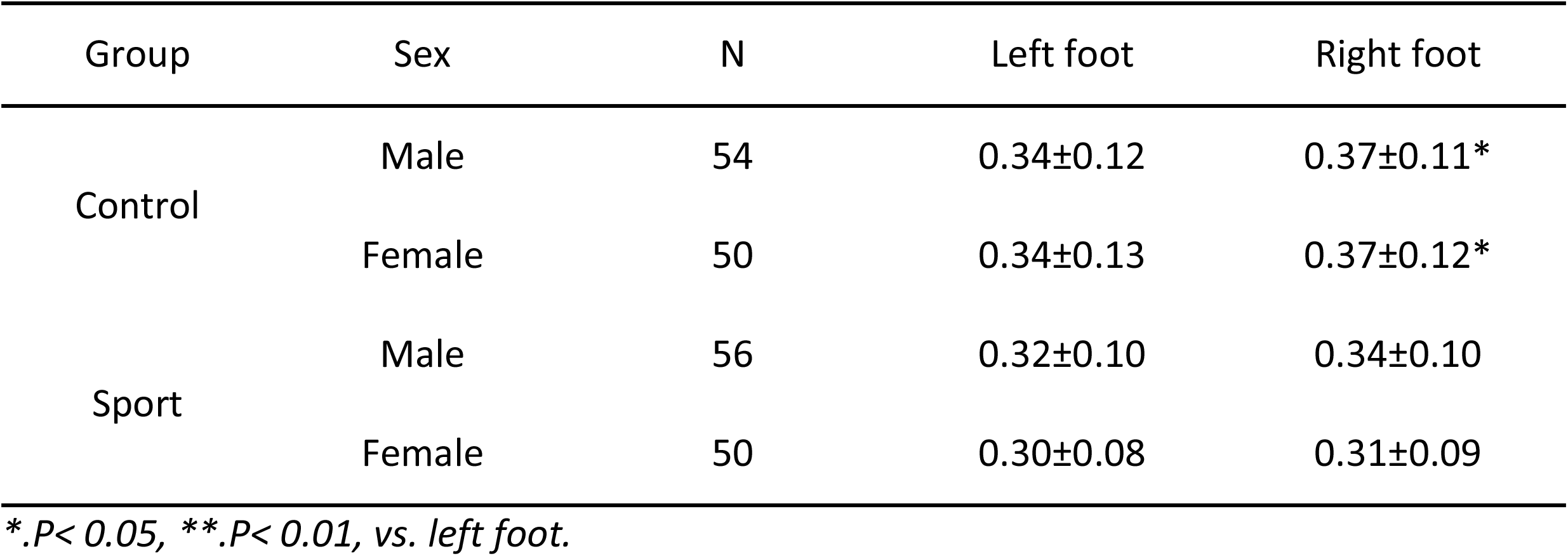
Comparison of CSI between left foot and right foot of the subjects

## Discussion

We analyzed CSI, SAI and other indicators of two-dimensional static footprints in this study to ensure the reliability of the two testers who measured the indicators of two-dimensional static footprints and X-ray photos. ICCs across the same-subject repeated measures trials were calculated for each of the two examiners (intra-rater) and between the two examiners (inter-rater). The differences between single measuring ICC and average measuring ICC depend on the research need. Single measuring ICC was the indicator of reliability of a single evaluator. Average measuring ICC was the correlation coefficient of the mean score or sum of multiple evaluators. In the experiment, average measuring ICC was applied since two testers copied the footprints, so interpretation of the ICCs was conducted in accordance with Portney and Watkins^13^. In Table 2, average measuring ICC of CSI is 0.976, of SAI is 0.975, which is similar to the results of Queen^12^ with the average measuring ICC of CSI and SAI as 0.961 and 0.962. It proves that the two testers are of high reliability. In addition, two-dimensional static footprint indicators, namely CSI and SAI, have high reliability *(CSI>0.800)*. Calcaneal inclination angle, calcaneal–first metatarsal angle and the ratio of arch height to arch length are universally accepted indicators in medial longitudinal arch evaluation. Although previous studies have proved the reliability of the above three indicators, different machine and software have been applied in the present study, therefore, we also evaluate the reliability of the two testers who measured the indicators of X-ray photos. Table 3 shows good reliability of the above three indicators. Indicators of the testing will not be affected through days in a short time. We focus on comparing the differences between athletes of high level and normal university students with Two-Dimensional Static Footprint. Moreover, we did not find measurement between days in our references^14^.

Two-dimensional static footprint analysis is one of the methods of arch evaluation. The traditional arch evaluation methods include the “Three Lines Method” and the “Ratio Method” (Figure 2). The “Three Lines Method” is defective, which causes large measuring error and makes it difficult to quantify the measurements. “The Ratio Method” is of higher accuracy with the ability to perform quantitative description, but it lacks strict definition on how to measure and draw. CSI and SAI, as indicators of two-dimensional static footprint analysis, have been used worldwide^4,12,14^. Since there exists difference in the foot shape among people of different races, gender and occupation, both athletes and normal people are selected as sport group and control group. As a whole, CSI, SAI and the indicators of the X-ray photos have a high relative correlation, which proves the validity of CSI and SAI in the evaluation of the relative height of the medial longitudinal arch among athletes and normal people. CSI and SAI are inversely correlated with calcaneal inclination angle and the ratio of height to length of the medial longitudinal arch and positively correlated with calcaneal-first metatarsal angle. This indicates, in arch evaluation, that the higher the relative height of the medial longitudinal arch is, the larger calcaneal inclination angle and the ratio of height to length of medial longitudinal arch are and the smaller calcaneal-first metatarsal angle, CSI and SAI are. It deserves attention that the correlation between either CSI or SAI and indicators of X-ray photos of the control group is higher than that of the sport group. From the anatomical point of view, the foot shape of the track and field athletes changes to some extent due to long-term exercise training, though there is rarely or no training specific for plantar muscle. Because the validity of testers on CSI and SAI in medial longitudinal arch evaluation are similar to each other and the sample size is great in Experiment Two, it’s redundant to measure both CSI and SAI. Based on the report that CSI enjoys a higher reliability than SAI in flatfoot determination, we chose CSI as the indicator to evaluate medial longitudinal arch.

Firstly, we compared CSI between athletes and college students in Experiment Two. We found that athletes’ mean CSI is smaller than that of college students, and female athletes’ CSI presented significant difference from that of the control group. It proved that the medial longitudinal arch of athletes is higher than that of college students, which is possibly due to exercise training. Twenty-one asymptomatic subjects (42 feet) completed a 4-week Short Foot Exercise training program. Compared with the data before the program, the amount of navicular drops decreased, and the arch height index increased, that is to say, this study offers preliminary evidence to suggest that Short Foot Exercise training may have value in statically and dynamically supporting the medial longitudinal arch^15^. Due to long-term exercise training, the relative height of the medial longitudinal arch of the track and field athletes is higher than that of normal people. Though there is rarely or no training specific for plantar muscle, quadratus plantae, lumbrical, flexor hallucis brevis and flexor digitorum brevis are used when the athletes are taking exercises to train their larger muscle groups.

In some parts of the world, the height of medial longitudinal arch is included in selection measures for athletes, that is, people with relatively higher medial longitudinal arch due to genetic inheritance are selected as athletes. The reason lies in that the relative height of medial longitudinal arch plays an important role in foot bio-mechanics when people are doing exercises with lower limbs. Researches based on bio-mechanics have proved that the heel valgus will take place more seriously and more quickly among runners with low arch foot^16^, which requires more control either initiative or passive over foot and the medial ankle, and the relevant ligaments and tendons will bear greater pressure accordingly. This may be helping to explain the advantage of athletes with relatively higher arch in running and jumping. Female athletes’ medial longitudinal arches are significantly higher than those of female college students. Although this cannot prove that the relative height of the medial longitudinal arch is directly related to running, jumping and other athletic abilities, it’s enough to explain that the relative height of medial longitudinal arch is an important indicator in track and field.

The toe function is of vital importance to balance control as well as change of body movements^17^. Length measurement of footprint is related to foot length, so it is common to apply the ratio of each measurement of footprint to foot length in foot morphology comparison^18^. As a whole, there is no apparent difference in toe length between athletes and college students. Long-term exercise training has helped to develop plantar muscle including quadratus plantae and lumbrical, but it has no effect on the relative toe length among high level athletes. Footprint length is related to body height^19^, and it is common to estimate body height from footprint length in forensic medicine.

Multiple studies have proved difference in foot morphology between bilateral feet^20–22^, but there are few studies focused on the asymmetry of bilateral medial longitudinal arch, which is one of the most important morphological structures of foot. Table 6 shows that there is no asymmetry of bilateral medial longitudinal arch among athletes, however, asymmetry of bilateral medial longitudinal arch does exist among normal people. Based on the fact that the shape of medial longitudinal arch is associated with genetic inheritance^10^, the asymmetry of bilateral medial longitudinal arch among normal college students is probably congenital. In sport group, however, the basic training movements in track and field like walking, running and jumping require cooperation between bilateral legs. Bilateral lower limbs move simultaneously or alternately, which ensures the same training on both dominant foot and nondominant foot. This helps to explain why there is no asymmetry of bilateral medial longitudinal arch among athletes.

## Perspectives

The medial longitudinal arch is one of the most important variables in studying footprint. We prove the qualification of CSI and SAI in medial longitudinal arch evaluation and explain that the relative height of medial longitudinal arch is an important indicator in track and field. However, the relative height of the medial longitudinal arch of the right foot is lower than that of the left foot among college students. We hope that the validity of two-dimensional static footprint analysis in medial longitudinal arch evaluation as well as the characteristics of athletes’ footprints was explored to provide a basis for the evaluation and selection of athletes.

## Acknowledgments

We appreciate the time and effort of the participants in this study. The work was supported by Youth Program of National Natural Science Foundation of China (Grant No. 81902298, 81901430).

## Contributors

Yu ZS, Yuan Y and Zhang LL designed this study. Zhang LL, Yu DL, Lei L, Gao YW and Dong JJ performed experiments. Yu DL and Zhang LL analyzed experimental data. Zhang LL and Yu DL were responsible for manuscript writing. Yuan Y revised the manuscript. All authors approved the final version of this manuscript.

## Patient consent

Obtained.

## REFERENCES

1 Reel S, Rouse S, Vernon W, et al. Reliability of a two-dimensional footprint measurement approach. Science and Justice 2010; 50(3): 113–8.

2 Reischl U, Nandikolla V, Colby C, et al. A case study of Chinese bound feet: application of footprint analysis. Collegium antropologicum 2008; 32(2): 629–32.

3 El O, Akcali O, Kosay C, et al. Flexible flatfoot and related factors in primary school children: a report of a screening study. Rheumatology international 2006; 26(11): 1050–3.

4 Reel S, Rouse S, Obe W V, et al. Estimation of stature from static and dynamic footprints. Forensic science international 2012; 219(1-3): 283. e1-5.

5 Chen K C, Yeh C J, Kuo J F, et al. Footprint analysis of flatfoot in preschool-aged children. European journal of pediatrics 2011; 170(5): 611–7.

6 Villarroya M A, Esquivel J M, Tomás C, et al. Assessment of the medial longitudinal arch in children and adolescents with obesity: footprints and radiographic study. European journal of pediatrics 2009; 168(5): 559–67.

7 Chang J H, Wang S H, Kuo C L, et al. Prevalence of flexible flatfoot in Taiwanese school-aged children in relation to obesity, gender, and age. European journal of pediatrics 2010; 169(4): 447–52.

8 Lin C J, Lai K A, Kuan T S, et al. Correlating factors and clinical significance of flexible flatfoot in preschool children. Journal of pediatric orthopaedics 2001; 21(3): 378–82.

9 Garcia-Rodríguez A, Martin-Jiménez F, Carnero-Varo M, et al. Flexible flat feet in children: a real problem?. Pediatrics 1999; 103(6): e84.

10 Kanatli U, Yetkin H, Cila E. Footprint and radiographic analysis of the feet. Journal of Pediatric Orthopaedics 2001; 21(2): 225–8.

11 Pauk J, Ezerskiy V, Raso J V, et al. Epidemiologic factors affecting plantar arch development in children with flat feet. Journal of the American Podiatric Medical Association 2012; 102(2): 114–21.

12 Queen R M, Mall N A, Hardaker W M, et al. Describing the medial longitudinal arch using footprint indices and a clinical grading system. Foot & ankle international 2007; 28(4): 45662.

13 Evans A M, Rome K, Peet L. The foot posture index, ankle lunge test, Beighton scale and the lower limb assessment score in healthy children: a reliability study. Journal of foot and ankle research 2012;, 5(1): 1.

14 Yalcin N, Esen E, Kanatli U, et al. Evaluation of the medial longitudinal arch: a comparison between the dynamic plantar pressure measurement system and radiographic analysis. Acta Orthop Traumatol Turc 2010; 44(3): 241–5.

15 Mulligan E P, Cook P G. Effect of plantar intrinsic muscle training on medial longitudinal arch morphology and dynamic function. Musculoskeletal Science & Practice 2013; 18(5): 425–30.

16 Williams Iii D S, McClay I S, Hamill J. Arch structure and injury patterns in runners. Clinical biomechanics 2001; 16(4): 341–7.

17 Kulthanan T, Techakampuch S, Donphongam N. A study of footprints in athletes and non-athletic people. J Med Assoc Thai 2004; 87(7): 788–93.

18 Mauch M, Mickle K J, Munro B J, et al. Do the feet of German and Australian children differ in structure? Implications for children’s shoe design. Ergonomics 2008; 51(4): 527–39.

19 Bosch K, Gerβ J, Rosenbaum D. Development of healthy children’s feet—nine-year results of a longitudinal investigation of plantar loading patterns. Gait & posture 2010; 32(4): 564–71.

20 Krishan K. Estimation of stature from footprint and foot outline dimensions in Gujjars of North India. Forensic science international 2008; 175(2-3): 93–101.

21 Zeybek G, Ergur I, Demiroglu Z. Stature and gender estimation using foot measurements. Forensic Science Internationa 2008; 181(1-3): 54. e1-5.

22 Domjanic J, Fieder M, Seidler H, et al. Geometric morphometric footprint analysis of young women. Journal of foot and ankle research 2013; 6(1): 27.

